# Differential restriction of chikungunya virus in primary human cardiac endothelial cells occurs at multiple steps in the viral life cycle

**DOI:** 10.1101/2024.09.13.612809

**Authors:** Sophie N. Spector, Maria G. Noval, Kenneth A. Stapleford

**Affiliations:** Department of Microbiology, New York University Grossman School of Medicine, New York, NY

## Abstract

Arthropod-borne viruses (arboviruses) constitute a significant ongoing public health threat, as the mechanisms of pathogenesis remain incompletely understood. Cardiovascular symptomatology is emerging as an important manifestation of arboviral infection. We have recently studied the cardiac tropism and mechanisms implicated in cardiac damage in mice for the alphavirus chikungunya virus (CHIKV), and we therefore sought to evaluate the cardiac tropism of other emerging alphaviruses and arboviruses. Using human primary cardiac cells, we found that arboviruses from diverse viral families were able to replicate within these cells. Interestingly, we noted that while the closely related alphavirus Mayaro virus (MAYV) could replicate to high titers in primary human cardiac microvascular endothelial cells, pulmonary, and brain endothelial cells, the Indian Ocean Lineage of CHIKV (CHIKV-IOL) was completely restricted in all endothelial cells tested. Upon further investigation, we discovered that this restriction occurs at both entry and egress stages. Additionally, we observed that compared to CHIKV, MAYV may antagonize or evade the innate immune response more efficiently in human cardiac endothelial cells to increase infection. Overall, this study explores the tropism of arboviruses in human primary cardiac cells and characterizes the strain-specific restriction of CHIKV-IOL in human endothelial cells. Further work is needed to understand how the differential restriction of alphaviruses in human endothelial cells impacts pathogenesis in a living model, as well as the specific host factors responsible.

**Author Summary:** Mosquito-borne viruses, such as those within the alphavirus genus, are an ongoing concern to human health globally. While we have recently begun to explore the mechanisms of CHIKV-induced pathology in the heart, little is known about other arboviruses and alphaviruses related to CHIKV. Here, we identified that multiple cardiac cell types are susceptible to infection by several arboviruses important to public health. Specifically, we noted differences in how two related viruses, CHIKV and MAYV, infect endothelial cells from multiple origins. This work highlights the potential for other emerging arboviruses, in addition to CHIKV, to directly infect cardiac tissue. Moreover, it emphasizes the ability for MAYV, an emerging virus that is less studied, to infect various endothelial cell types to high titers, suggesting the need for further research on MAYV pathogenesis. Finally, this research identifies differences in infection between individual strains of CHIKV, suggesting a finely tuned mechanism of restriction in endothelial cells that can be further explored, as well as the need to study use different viral strains for future alphavirus research.

## Introduction

Arboviruses include human pathogens important to global public health. Despite their increasing epidemic potential due to climate change and globalization, the mechanism of pathogenesis of many emerging arboviruses are not yet fully understood. With the increased circulation of these viruses in higher-income countries with improved health surveillance, data supporting a direct link between arboviral infections and severe manifestations have started to emerge. In particular, cardiovascular complications are a concerning outcome reported after infection with dengue virus, Zika virus, and chikungunya virus (CHIKV), among other arboviruses [1–7]. Investigations of CHIKV have identified viral antigen and viral RNA in cardiac tissue of individuals that succumbed during the acute and post-acute phases of the infection, supporting a direct infection of the tissue [8, 9]. Given these findings, the potential for other arboviruses to cause cardiac damage through direct infection of the heart is of clinical concern; arboviral infections are underreported worldwide, and therefore, cardiovascular manifestations in patients may be poorly surveilled.

CHIKV is an alphavirus that is endemic in Africa, India, Southeast Asia, and South America. Using mouse models along with primary human cardiac cells, we have recently demonstrated a direct link between heart infection and cardiac tissue inflammation [10]. We identified cardiac fibroblasts as the main target of CHIKV infection in the heart of mice [10]. Our findings on CHIKV raise questions about whether related alphaviruses and other arboviruses, many of which have the potential to emerge in new environment, can infect cardiac tissue. Other arthritogenic alphaviruses in the Semliki Forest virus complex include Mayaro virus (MAYV) and Ross River virus (RRV), which are less commonly studied despite being closely related to CHIKV. MAYV is of particular interest as it is responsible for small human outbreaks in South and Central America and has a similar clinical presentation as CHIKV [7]. Although many cases are likely misdiagnosed, confirmed cases of MAYV have been steadily increasing, and it is thought that the virus has the potential to adapt to urban vectors [11]. Interestingly, MAYV RNA has been recently identified in different regions of the heart tissue in rhesus macaques 10 days post-infection [12].

Here, we evaluated the susceptibility of different primary human cardiac cell types to several arboviruses of public health concern. We found that arboviruses from diverse families have the potential to infect human cardiac cells *in vitro*. Interestingly, the CHIKV Indian Ocean Lineage (CHIKV-IOL) is uniquely restricted in human cardiac microvascular endothelial cells (hCMECs). This restriction was also observed in human pulmonary and brain microvascular endothelial cells. However, in line with previous studies in endothelial cells using other CHIKV strains, we found that this cell type specific restriction was strain dependent and specific to CHIKV-IOL [13–15]. We found that CHIKV-IOL infection was restricted at both entry and egress. Interestingly, we can rescue CHIKV-IOL restriction by prolonged treatment of hCMECs with the JAK/STAT inhibitor ruxolitinib for up to four days, suggesting a role of the innate immune response in this restriction. Investigating innate immune responses after infection exposed differences in interferon-stimulated genes (ISG) upregulation between alphaviruses, suggesting that MAYV may antagonize or evade IFN-I responses in endothelial cells more efficiently compared to CHIKV-IOL.

Overall, this work suggests cell type specific restrictions between alphaviruses, while also exposing tropism for human cardiac cells across genetically diverse arboviruses. Moreover, we find an endothelial cell-specific mechanism of antiviral restriction of CHIKV-IOL, which other CHIKV strains and alphaviruses may be able to antagonize or otherwise bypass more efficiently. More broadly, we found that the differential restriction of alphaviruses in cardiac endothelial cells model phenotypes across various human endothelial cell types, signifying possible repercussions for differences in pathogenesis across organs. Although further work is needed to characterize the specific molecular mechanism of restriction, and the characteristics of MAYV that allow it to efficiently avoid restriction, this study seeks to characterize differences in alphavirus restriction that may impact pathogenesis both in the heart and across endothelial cells.

## Methods

### Biosafety

All work with the Indian Ocean Lineage of chikungunya virus (CHIKV-IOL) was completed under Biosafety Level 3 (BSL3) conditions at the NYU Grossman School of Medicine.

### Cells

Vero cells (ATCC CCL-81) were grown in Dulbecco’s Modified Eagle Medium (DMEM, Corning) supplemented with 10% newborn calf serum (NBCS, Sigma). BHK-21 cells (ATCC CCL-10) and BSR-T7 cells (a gift from Dr. Steven Whitehead at the National Institutes of Health (NIH) [43] were grown in DMEM supplemented with 10% fetal bovine serum (FBS, Atlanta Biologicals), 1% nonessential amino acids (NEAA, Corning), and 10 mM HEPES (Invitrogen) with the addition of 1 mg/ml geneticin every other passage to maintain selection. Human primary cardiac cells (PromoCell) were grown following the company’s recommendations, and passaged using the DetachKit (PromoCell, C-41200).

Human cardiac fibroblasts (hCF, C-12375) were grown in Fibroblast Growth Medium 3 (PromoCell, C-23025), human cardiac myocytes (hCM, PromoCell, C-12810) were grown in Myocyte Growth Media (PromoCell, C-22070), human aortic smooth muscles (hAoSMC, PromoCell, C-12533) were grown in Smooth Muscle Cell Growth Medium 2 (PromoCell, C-22062), and human cardiac microvascular endothelial cells (hCMEC, PromoCell, C-12285) were grown in Endothelial Cell Growth Media MV (PromoCell, C-22020). One donor was used for each human primary cardiac cell type. Primary cells were used for experiments up to passage 11. All primary cell media was supplemented with 1% penicillin-streptomycin (Pen/strep, Cellgro, 30-002-CI). Immortalized human brain microvascular endothelial cells (hBMEC) and immortalized human pulmonary microvascular endothelial cells (hPMEC) [44, 45], gifts from Dr. Ana Rodriguez at the NYU Grossman School of Medicine were used between P11 and P15 for all experiments, and grown in ECM medium supplemented with 5% FBS, 1% endothelial cell growth supplement, and 10 mg/ml pen/strep (ScienCell, 1001). All cells were maintained at 37°C in 5% CO_2_ and confirmed mycoplasma-free using the Lookout Mycoplasma PCR detection kit (Sigma-Aldrich).

### Viruses

[21]Wild-type CHIKV-IOL (Strain 06-049, AM258994) [46], CHIKV-IOL-ZsGreen, CHIKV-Caribbean (LN898104.1)[47], CHIKV-Asian (Strain Mal06; EU703759.1, a gift from Dr. Scott Weaver at the University of Texas Medical Branch (UTMB)) [48], CHIKV-181/25 (L37661, a gift from Dr. Laurie Silva at the University of Pittsburgh), SINV (NC_001547.1, a gift from Dr. Benjamin tenOever at the NYU Grossman School of Medicine) were generated from *in vitro* transcribed RNAs derived from infectious clones. Briefly, 10 μg of each infectious clone plasmid was linearized overnight with NotI (CHIKV plasmids) or XhoI (SINV plasmid) restriction endonucleases (Invitrogen), and then purified by phenol:chloroform extraction and ethanol precipitation. RNA was *in vitro* transcribed using the SP6 mMessage mMachine kit (Invitrogen) following the manufacturer’s instructions, and purified by phenol:chloroform extraction and ethanol precipitation. BHK-21 cells (10^7^ cells/ml) were electroporated with 10 μg of *in vitro* transcribed RNA by 1 pulse of 1,200 V, 25 Ω, and infinite resistance. Electroporated cells were added to a T25 flask in complete media. After incubation at 37°C for 48-72 hours, the passage 0 (p0) supernatant was centrifuged at 1,200 rpm for 5 minutes. A passage 1 (p1) working stock was generated by infecting BHK-21 cells with p0 virus, collecting supernatant after incubation at 37°C for 48-72 hours, and centrifuging at 1,200 rpm for 5 minutes. Ultracentrifugation was used to generate purified virus stocks. Viruses were pelleted over a 20% sucrose cushion by centrifugation at 25,000 rpm for 4 hours. Purified virus particles were resuspended in media consisting of DMEM containing 2% FBS.

Mayaro virus (TRVL4675) was a gift from Drs. Gonzalo Moratorio and Alvaro Fajardo at the Institut Pastuer, Montevideo, Uruguay. Ross River virus (Strain T-48, NR-51457) and La Crosse virus (AF528165, NR-540) were both obtained from BEI resources. MAYV, RRV, and LACV were passaged in Vero cells to generate a working stock as above. RVFV MP-12 was generated by transfecting 2 μg of plasmids encoding the S, M, and L segments, gifts from Dr. Shinji Makino at UTMB, into BSR-T7 cells using the LT-1 transfection reagent (Mirus), and passaged to generate a P1 stock as above. The Zika virus (Brazilian strain) infectious clone plasmid, a gift from Dr. Alexander Pletnev at the NIH, was transfected into 293T cells in a 6-well plate using the lipofectamine 2000 transfection reagent (Thermo Fisher), and passaged in Vero cells to generate working stock as above. Infectious virus titers were quantified by plaque assay for all stocks as described below

### Plaque assay

Infectious virus production was quantified by plaque assay. Virus samples were diluted 10-fold in DMEM and added to a monolayer of Vero cells for 1 h at 37 °C. After incubation, cells were overlaid with 0.8% agarose in DMEM supplemented with 2% NBCS and 1% antibiotic-antimycotic (Gibco). Cells were incubated at 37 °C for 48 h (MAYV), 72 h (CHIKV, LACV, or RVFV MP-12), or 96 h (ZIKV) and fixed with 4% formalin. The agarose plugs were removed, and plaques were visualized by crystal violet staining. Viral titers were determined by counting plaques in the highest countable dilution.

### Primary human cardiac cell and endothelial cell growth curves

For growth curves in cardiac cells, human primary cells (hCF, hAoSMC, hCM, hCMEC) were seeded in a 24-well plate at a density of 50,000-90,000 cells/well depending on cell type. 24 h later, RVFV MP-12, LACV, ZIKV, and MAYV were diluted in DMEM to a MOI of 0.1. After removing complete media, virus was added to the cells and incubated for 1 h at 37°C. After incubation, the inoculum was removed and cells were gently washed with PBS twice, and complete media specific to each cell type was replenished. At 0, 6, 24, 48, and 72 hours post infection (hpi), half of the supernatant was collected to quantify infectious particle production by plaque assay as described above, and the plate was replenished with complete media.

For assays comparing endothelial cell types, CHIKV strains, and other alphaviruses, hCMECs, hBMECs, and hPMECs were seeded in a 96-well plate at a density of 10,000-15,000 cells/well. 24 h later, each virus was diluted in DMEM to a MOI of 0.1 and added to the cells after removing media. After 1 h of incubation at 37°C, inoculum was removed, the cells were gently washed once with PBS, and complete media specific to each cell type was added. At 48 hpi, the supernatant was collected to quantify infectious particle production by plaque assay as described above. For all virus infection experiments, virus inputs were verified by plaque assay.

### Immunofluorescence staining

48 hours post infection, cells were fixed with 4% paraformaldehyde for 1 hour. Cells were then washed with Perm/Wash (BD Biosciences) and incubated with 0.25% TX-100 for 10 minutes. Cells were incubated in blocking buffer (0.2% BSA, 0.05% saponin, in PBS) for 1 hour followed by the addition of a rabbit anti-CHIKV capsid antibody (a gift from Dr. Andres Merits at the University of Tartu, Estonia) for 2 hours. Wells were washed three times with Perm/Wash and incubated with anti-rabbit-Alexa555 secondary antibody and DAPI for 1 hour followed by extensive washing. Images were acquired using the CellInsight CX7 imaging platform.

### CellInsight CX7 quantification

To quantify the number of infected cells after alphavirus infection, cells were fixed with 4% formalin for 1 h at room temperature. Cells that were infected with CHIKV-IOL-ZsGreen were stained with 4’,6-diamidino-2-phenylindole (DAPI, Thermo Scientific) diluted 1:1000 in PBS and incubated for 1 h at room temperature. A CellInsight CX7 high-content microscope (Thermo-Scientific) was used to quantify the percent of infected cells in each well by quantifying the number of ZsGreen positive cells compared to the total number of cells measured by DAPI.

### Fusion-from-without-assay

HCMECs were seeded in 96-well flat transparent black plates (Corning) at a density of 12,000 cells/well. The next day, media was removed, cells washed once with binding buffer (Endothelial Cell Growth Media MV supplemented with 0.2% BSA, 10 mM HEPES, and 20 mM NH_4_Cl) and incubated in 100 μL of binding buffer for 90 minutes at 4°C. After incubating in binding buffer, CHIKV-IOL and MAYV stocks were diluted in binding buffer to a MOI of 5, buffer was removed, and virus dilutions were added to cells for 1 h at 4°C. After incubation, the virus inoculum was removed while the plate remained on ice. Fusion buffers were made by supplementing Endothelial Cell Growth Media MV with 0.2% BSA, 10 mM HEPES, and 30 mM succinic acid adjusting the pH to either 7.5 or 5.2 with NaOH, and sterile filtering prior to use. Pre-warmed fusion buffer at pH 7.5 or 5.2 was then added to the cells, and incubated at 37°C for 2 minutes. The fusion buffer was promptly removed, washed once with complete endothelial cell growth media MV, and replenished with complete media. At 24 and 48 hpi, a portion of supernatant was collected to quantify infectious virus by plaque assay. At 48 hpi, the plate was fixed with 4% formalin, washed three times with PBS, and stained with DAPI for high-content CX7 microscopy as described above.

### Ruxolitinib treatment assays

For 48 h pre-treatment infection assays, hCMECs were seeded in 96-well flat transparent black plates at a density of 12,000 cells/well. The next day, cells were treated with 5 μM ruxolitinib (Invitrogen, INCB018424) diluted in complete endothelial cell media or mock-treated. After 24 h, the treated condition was replaced with new media with fresh ruxolitinib, to mitigate the effects of degradation, and treated for another 24 h. After 48 h total of ruxolitinib or mock pre-treatment, cells were then infected with CHIKV-ZsGreen diluted in DMEM to a MOI of 0.1, or mock-infected with DMEM, and incubated at 37°C for 1 h. After incubation, the inoculum was removed, cells were washed once with PBS, and complete Endothelial Cell Growth Media MV was added, with 5 μM ruxolitinib supplemented in each ruxolitinib condition and cells were then incubated at 37°C. At 0, 1, 2, and 4 days post-infection half of the supernatant was collected to quantify infectious particles by plaque assay and replaced with complete media supplemented with 5 μM ruxolitinib in the treatment condition. At 5 dpi, the plate was fixed in 4% formalin for 1 h, washed with PBS three times, and stained with DAPI as described above. Infected cells were then quantified by high-content microscopy on the CX7 microscope as described above.

For assessment of host-proteins after ruxolitinib treatment, hCMECs were seeded in 12-well flat transparent plates at a density 90,000 cells/well. The next day, the 6-day pre-treatment condition was treated with 5 μM ruxolitinib for 48 h, while the 4-day pre-treatment and mock-condition cells were left in complete media. The 6-day pre-treatment ruxolitinib media was replaced with new media with fresh ruxolitinib after 48 h, while the 4-day ruxolitinib pre-treatment was started. After another 48 h, the 5 uM ruxolitinib media was replaced for both conditions, and the 2-day pre-treatment was started. After a final 48 h, the cell monolayer was washed once with PBS and then harvested in RIPA buffer (50 mM Tris pH 8.0, 150 mM NaCl, 1% Tx-100, 0.5% sodium deoxycholate, 0.1% SDS, and halt protease inhibitor cocktail (Thermo)) for western blotting as described below.

### Single-cell analysis of *Tabula sapiens*

Single-cell heart and lung data from *Tabula sapiens* [26] was downloaded from https://figshare.com/projects/Tabula_Sapiens/100973. Data was analyzed in R (version 4.3.2) using the Seurat (version 5.0.1) [49] and SeuratDisk R packages. Data was normalized and clustered, maintaining the original cluster annotations from *Tabula Sapiens*. A dot plot representing normalized expression of cell markers was generated using the “DotPlot” function.

### Recombinant IFN-β Treatment

HCMECs, hPMECs, hBMECs, and hCFs were seeded in a 12-well plate at 80,000–90,000 cells per well. The following day, one well of each cell type was treated with recombinant human IFNβ (Millipore Sigma, #IF014) diluted to 100 U/ml in appropriate media, while another well was mock-treated. 24 h later, cells were washed once with PBS and then collected in RIPA buffer for analysis by western blotting as described below.

### Western blotting

Confluent cells in a 12-well plate were washed once with PBS and then collected in RIPA buffer with 1x Halt protease inhibitor cocktail (Thermo). After lysis, sample was mixed in equal proportion with 2X Laemmli buffer containing 10% β-mercaptoethanol, and boiled at 95°C for 10 mins. Debris was removed by centrifugation at 10,000 x g for 5 mins. Samples were separated by SDS-PAGE on a 12% acrylamide gel and transferred to a hydrophobic polyvinylidene fluoride (PVDF) membrane (Immobilon). Membranes were then incubated in blocking buffer (1X Tris buffered saline (TBS; 20 mM Tris-HCl, pH 7.4, 150 mM NaCl, 1 mM EDTA, 1% TX-100), 0.1% Tween-20%, and 5% dry milk) overnight at 4°C. The following day, blots were incubated in primary antibodies including Actin (MA5-11869, ThermoFisher), IFITM2 (12769-1-AP, Proteintech), and IFITM3 (PA5-11274, ThermoFisher) for 2-3 h at room temperature and washed with 1X TBST. Then, membranes were incubated in anti-mouse or anti-rabbit IgG HRP secondary antibodies (Invitrogen) for 1 h at room temperature, and washed in 1X TBST again. Finally, blots were developed using a SuperSignal Weser Pico plus chemiluminescence substrate kit (Thermo) and imaged on a ChemiDoc MP imaging system (Bio-Rad). Images were analyzed and bands quantified using Image Lab (Bio-Rad). Relative expression is quantified as the adjusted band volume of target gene divided by the adjusted band volume of the actin loading control. Protein loading was further verified by staining membranes with Coomassie blue.

### Type I interferon response assays and poly(I:C) transfection

HCMECs were seeded in a 24-well plate at a density of 45,000 cells/well. The next day, cells were transfected with 312 ng of high molecular weight poly(I:C) (Invitrogen) or mock-transfected using TransIT-mRNA Transfection Kit (Mirus Bio) and incubated for 2 h at 37°C. Afterwards, cells were mock-infected with DMEM, infected with CHIKV-IOL, CHIKV-181/25, or MAYV at a MOI of 0.1 diluted in DMEM, or infected with heat-inactivated stocks of each virus. For this assay, each virus stock was produced in low-passage BHK-21 cells, and ultra-purified over a 20% sucrose cushion by ultracentrifugation at 25,0000 x g for 2 hours at 4°C. Heat-inactivated virus was produced by incubating virus stocks at 56°C for 4 h. Infected cells were then incubated at 37°C for 1 h. After 3 h poly(I:C) or mock-transfection treatment, or 1 h virus incubation, cells were washed with PBS twice, and complete endothelial cell media was added. Cells were then returned to incubate at 37°C. At 0 and 24 hpi, supernatant was collected to quantify infectious particle production by plaque assay as previously described. Additionally, cells were washed once with PBS, and cell monolayers were collected in 500 µL TRIzol^TM^ reagent (Invitrogen) for RNA extractions and RT-qPCR as described below. Initial viral dilutions were verified by plaque assay.

### RNA extraction and RT-qPCR

Relative post-infection *ISG15* and *CHIKV* RNA levels were evaluated by SYBR Green and TaqMan ^TM^ RT-qPCR, respectively. First, RNA extractions were performed using TRIzol^TM^ reagent following the manufacturer’s guidelines. Extracted RNA was resuspended in 400 µL of water. Initial viral dilutions and post-infection CHIKV samples were quantified using the TaqMan^TM^ RNA-to-C_T_ 1-step kit (Applied Biosystems) following the manufacturers guidelines in a total reaction volume of 25 µL/well. For TaqMan assays, a standard curve from 10 ng/μl to 1E-7 ng/μl of *in vitro* transcribed CHIKV viral RNA for each strain was generated as previously described [10]. The following primers and probes were used to amplify the nsP4 fragment of each CHIKV strain: CHIKV-IOL (fw:5–’TCACTCCCTGCTGGACTTGATAGA-3’, rv:5’-TTGACGAACAGAGTTAGGAACATACC-3’, and probe: (5′-(6-carboxyfluorescein)-AGGTACGCGCTTCAAGTTCGGCG-(black-holequencher)-3′), CHIKV 181/25 (fw:5–’TCACTCCCTGTTGGACTTGATAGA-3’, rv:5’-TTGACGAACAGAGTTAGGAACATACC-3’, and probe: 5′-(6-carboxyfluorescein)-AGGTACGCGCTTCAAGTTCGGCG-(black-holequencher)-3′. RT-qPCR was performed on QuantStudio 3 qPCR instrument using the following protocol: 48 °C for 30 min, followed by 10 min at 95 °C. Amplification was performed over 40 cycles as follows: 95 °C for 15 sec, 60 °C for 1 min.

For analysis by SYBR Green qPCR, cDNA was synthesized using the Maxima H minus-strand kit (Thermo) with random primers. The cDNA thermocycler protocol consisted of 25°C for 10 mins, 50°C for 30 mins, 85°C for 5 mins. cDNAs were then used with the SYBR Green Master Mix (Thermo) following the manufacturer’s guidelines, with a total reaction volume of 20 µL/well. Amplification on a QuantStudio 3 qPCR instrument was performed over 40 cycles as follows: 95°C for 15 s, and 60°C for 1 min. The melt curve was evaluated for each reaction, and data was collected using QuantStudio software v1.4**. Primers used: *ISG15* (fw:5–’TCCTGGTGAGGAATAACAAGGG-3’, and rv:5’-GTCAGCCAGAACAGGTCGTC-3’and *GAPDH* (fw:5–’GCAAATTTCCATGGCACCGT-3’, and rv:5’-GCCCCACTTGATTTTGGAGG-3’). The relative expression of *ISG15* to *GAPDH* was calculated as ΔΔCT, and expressed as fold-change normalized to mock-infected control. The ΔΔCT was calculated for each sample against the average ΔCT of the mock-infected condition within each time-point. All plates were run with technical duplicates and non-template controls.

### Statistics and data analysis

Statistical significance was given where p values were <0.05 using Prism Version 10.1.0. Specific tests are indicated in figure legends. All experiments were performed in either biological triplicate with technical triplicates or duplicates.

### Data availability

All data are available in this manuscript.

## Results

### Human primary cardiac cells are susceptible to diverse arboviruses *in vitro*

Cardiovascular manifestations are a rare but serious outcome reported after infection with several emerging arboviruses, including Zika virus (ZIKV) and chikungunya virus (CHIKV) [3–7]. In previous work, we have demonstrated that CHIKV can infect and replicate within mouse cardiac fibroblasts *in vivo* and human primary cardiac fibroblasts *in vitro* [10]. However, the question still remains as to whether this is specific to CHIKV or if other arboviruses can infect human cardiac cells. To address this question, we used human primary cardiac cells as a physiologically relevant model to evaluate whether arboviruses from genetically diverse viral families can infect heart cells *in vitro*. We selected four cell types that are known to compose the majority of heart tissue: cardiac fibroblasts (hCFs), cardiac myocytes (hCMs), microvascular endothelial cells (hCMECs), and smooth muscle cells of aortic origin (hAoSMCs) [16]. We then selected a panel of emerging arboviruses, including viruses with and without reported cardiac manifestations, to evaluate infection in primary cardiac cells. We selected two bunyaviruses, La Crosse virus (LACV) & Rift Valley Fever virus (RVFV, vaccine strain MP-12), a flavivirus, Zika virus (ZIKV), and an alphavirus closely related to CHIKV, Mayaro virus (MAYV).

We first evaluated the kinetics of infection in each primary cell type by seeding each primary cell type at a low passage, infecting with each virus at a MOI of 0.1, and collecting supernatant at 6, 24, 48, and 72 hours post infection (hpi) to quantify infectious virus by plaque assay (**Fig 1 A-D**). We found that each virus tested was able to infect, replicate and produce infectious particles within each primary human cardiac cell type, though with differences in the kinetics and magnitude of infectious virus production between viruses and between cell types. In particular, LACV and ZIKV peaked at a lower viral titer in cardiac myocytes than in any other cell types. LACV, an encephalitic bunyavirus, had lower titers than RVFV in cardiac myocytes and aortic smooth muscle cells, another bunyavirus with known cardiopulmonary symptomatology [17] while ZIKV infected all human cardiac cell types, in accordance with known cardiovascular manifestations of ZIKV infection in humans [3, 5]. Interestingly, the alphavirus MAYV reached the highest titers of all viruses in each cell type, with notably higher titers in cardiac microvascular endothelial cells than other viruses (**Fig 1D**). These results indicate that multiple arboviruses have the potential to infect human cardiac tissue.

**Figure 1.**
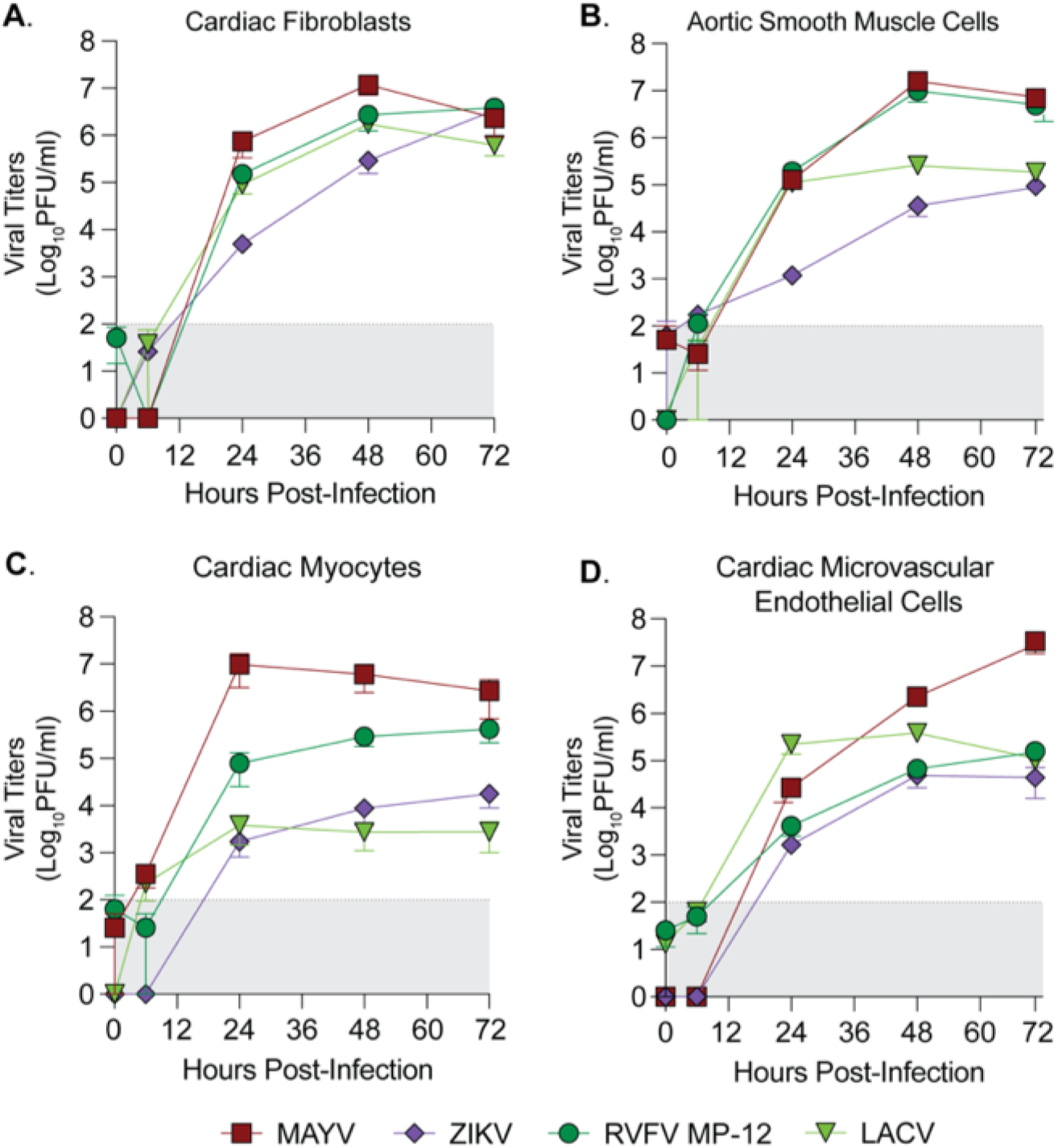
Human primary cardiac cells are susceptible to infection by diverse arboviruses. Human primary (**A**) cardiac fibroblasts, (**B**) aortic smooth muscle cells, (**C**) cardiac myocytes, and (**D**) cardiac microvascular endothelial cells were infected with MAYV, ZIKV, RVFV MP-12, or LACV at a MOI of 0.1. Supernatant was collected at 0, 6, 24, 48, and 72 hpi, and viral titers were quantified by plaque assay. Data points represent the mean of *n* = 2 independent trials with internal technical duplicates, with error bars representing the SEM. The limit of detection is indicated by the gray shaded area.

### The Indian Ocean Lineage of CHIKV is specifically restricted across human endothelial cell types

The fact that MAYV infected cardiac endothelial cells was interesting as we have previously shown that the Indian Ocean Lineage of CHIKV (CHIKV-IOL) was restricted in cardiac endothelial cells both *in vitro* and in immunocompetent WT C57BL/6 mice as measured by virus-cell colocalization analysis [10]. CHIKV and MAYV are closely related arthritogenic alphaviruses in the Semliki Forest virus complex which both utilize the Mxra8 receptor [18, 19]. Therefore, it was particularly interesting that MAYV could replicate to a high titer in primary hCMECs (**Fig 1D**), and by 72 hpi, we noted severe cytopathic effects (CPE) (data not shown). Given the similarity between CHIKV and MAYV, we sought to further characterize the differential restriction of these alphaviruses in human endothelial cells.

We first wondered if the restriction of CHIKV was specific to the CHIKV-IOL strain, or if this restriction was true in other CHIKV strains. To begin, we evaluated the growth of the Caribbean and Asian strains of CHIKV as well as the vaccine strain CHIKV-181/25 and CHIKV-IOL **(Fig 2A)**. We infected low passage hCMECs with each virus at a MOI of 0.1, and collected supernatants at 48 hpi to quantify infectious virus by plaque assay. We found that all other strains of CHIKV evaluated were able to productively infect hCMECs, while CHIKV-IOL was consistently undetectable at 48 hpi (**Fig 2A**).

**Figure 2.**
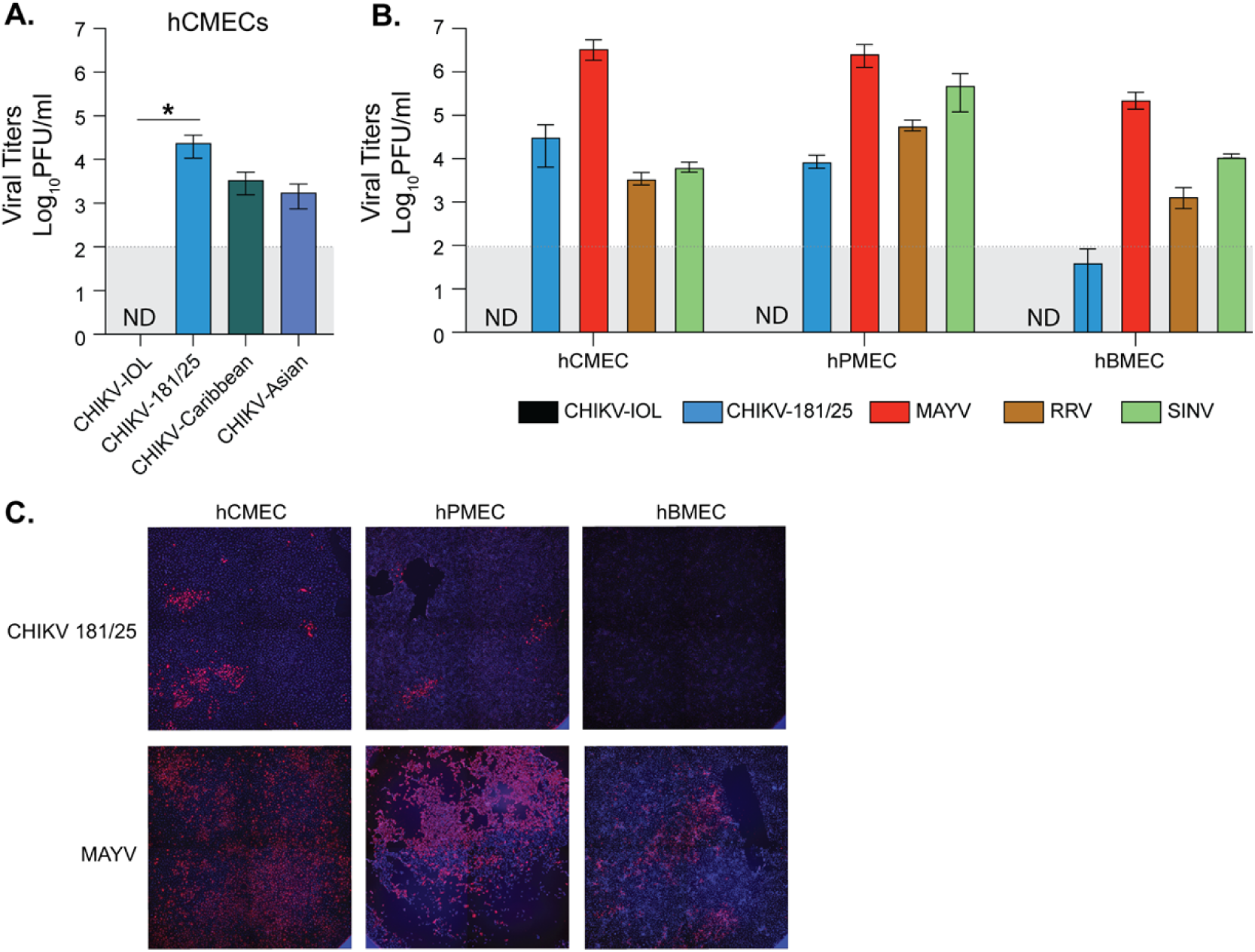
Differential growth of alphaviruses in endothelial cells. (**A**) hCMECs were infected with CHIKV-IOL, CHIKV-181/25, the CHIKV-Caribbean strain, or the CHIKV-Asian strain at an MOI of 0.1. Viral supernatants were collected at 48 hpi and infectious virus was quantified by plaque assay. **(B)** hCMECs, hPMECs, and hBMECs were infected with CHIKV-IOL, CHIKV-181/25, MAYV, RRV, or SINV at a MOI of 0.1. Supernatant was collected 48 hpi and viral titers were quantified via plaque assay. (**C**) At 48 hpi, cells were fixed and stained with an anti-CHIKV-capsid antibody and DAPI. Data represents the mean of at least *n* = 3 independent trials performed in technical triplicate, with error bars representing the SEM. The limit of detection (LOD) is indicated by the gray shaded area. Statistical significance was found via a Kruskal-Wallis test with Dunn’s multiple comparisons test (**B**), with p-values representing * p < 0.05. ND = Not detected.

Next, we questioned whether this differential restriction and growth could be expanded to different endothelial cells and alphaviruses. To address these questions, we infected hCMECs along with immortalized human brain microvascular endothelial cells (hBMECs) and immortalized human pulmonary microvascular endothelial cells (hPMECs) with MAYV, two strains of CHIKV – CHIKV-IOL and the vaccine strain CHIKV-181/25, Mxra8-dependent Ross river virus (RRV) of the SFV complex, as well as Sindbis virus (SINV), a more distantly related alphavirus in the western equine encephalitis complex [18, 19] – at a MOI of 0.1 **(Fig 2B and C**). We collected supernatants at 48 hpi and evaluated viral titers by plaque assay. We found that CHIKV-IOL was consistently restricted across human endothelial cell types, while MAYV reached significantly higher titers in hCMECs. Interestingly, the CHIKV-181/25 strain, adapted from a Southeast Asian human isolate [20], was able to infect cardiac and pulmonary endothelial cell types, although producing a lower viral titer than MAYV, yet was restricted in brain endothelial cells. These results suggest that this restriction may be specific to the CHIKV-IOL strain across endothelial cells. SINV and RRV both replicated in each endothelial cell line, although to much lower levels than MAYV. When we looked at the pattern of infected cells in each endothelial cell line between MAYV and CHIKV-181/25, we found that MAYV was able to spread throughout the cell monolayer over the course of infection while CHIKV-181/25 was found in distinct foci in hCMECs and hPMECs (**Fig 2C**). Taken together, these results suggest that CHIKV is restricted in a strain-specific manner across multiple human endothelial cell types, while MAYV replicates to higher titers with more robust spreading than other related CHIKV strains and alphaviruses.

### CHIKV-IOL is restricted at both early and late steps in virus life cycle

Given that CHIKV-IOL was restricted in multiple endothelial cell types, we next questioned where in the viral life cycle restriction was occurring. The fact that other CHIKV strains were able to both enter and productively replicate in hCMECs suggested that these cells do not lack the receptor or replication co-factors necessary for CHIKV entry and replication. To determine at which step in the alphavirus life cycle CHIKV-IOL was restricted, we performed a ‘fusion from without’ assay [21, 22] using our previously characterized CHIKV-IOL-ZsGreen reporter virus, which expresses a ZsGreen fluorescent protein only during active viral replication [21]. HCMECs were incubated with CHIKV-IOL-ZsGreen or MAYV at an MOI of 5 at 4°C in the presence of 20 mM NH_4_Cl to block endocytosis. Then, cells were exposed to a neutral (pH 7.5) or low pH buffer (pH 5.2) at 37°C for 2 minutes. In the low pH condition, the virus bypasses the endocytic pathway and releases viral contents directly through the plasma membrane. Therefore, if restriction is occurring after binding, but before replication, we would expect the ZsGreen reporter to be visible in the low pH condition. To evaluate infection, cells were fixed at 48 hpi with 4% formalin, stained with DAPI, and imaged by high-content microscopy.

In the neutral pH condition, we noted the restriction of CHIKV-IOL in hCMECs was reproducible even at a higher MOI of 5 and with a CHIKV-IOL-ZsGreen stock (**Fig 3B**). However, at a pH of 5.2 we saw an approximately 71-fold increase in CHIKV replication relative to the neutral pH condition, as measured by % of cells positive for ZsGreen (**Fig 3A and B**). Indeed, the percent of cells infected was significantly different from the mock-infected control in the low pH 5.2 condition, but not the pH 7.5 condition (**Fig 3A**). This finding suggests that inhibition of CHIKV-IOL is occurring at a post-binding but pre-replication step and can be partially recovered by bypassing endocytic entry. Notably, we did not see spreading by microscopy in the low pH condition, suggesting that infectious particles are not being produced, or that neighboring cells are still effectively restricting CHIKV-IOL (**Fig 3B**).

**Figure 3.**
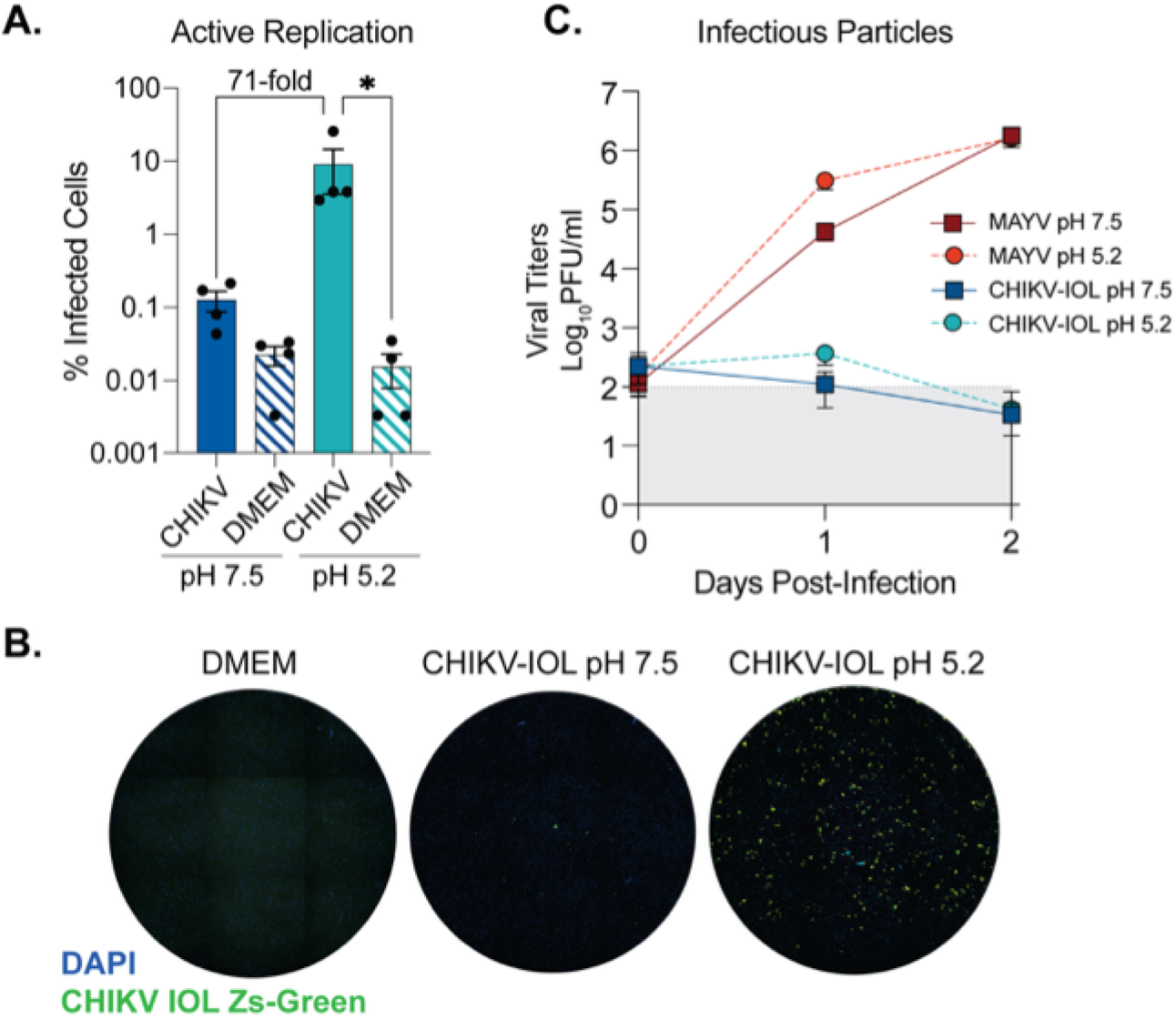
CHIKV IOL restriction occurs at viral entry and egress in cardiac endothelial cells. A fusion-from-without-assay was performed in hCMECs. CHIKV-IOL-ZsGreen or MAYV was bound to hCMECs at a MOI of 5 at 4°C for one hour. After incubation, buffer at a pH of 5.2 or 7.5 was added and incubated at 37°C for 3 mins, replaced with complete media and incubated for 2 days. Supernatant was collected at 0, 1, and 2 dpi, and cells were fixed and visualized by high-content microscopy 2 dpi. (**A**) CHIKV infection quantified as the percent of CHIKV-IOL-ZsGreen positive cells as compared to the total number of cell nuclei as indicated by DAPI staining by high-content microscopy. (**B**) Representative microscopy images showing uninfected (DMEM) and CHIKV-IOL-ZsGreen (green channel) infected cells at pH 5.2 and 7.5 with DAPI-stained cell nuclei (blue channel). (**C**) Viral titers from supernatant quantified via plaque assay. Data represents the mean of at least *n* = 3 independent trials in technical duplicate or triplicate, with error bars showing the SEM and the limit of detection indicated by the gray shaded area. Statistical significance was found via a Kruskal-Wallis test with Dunn’s multiple comparisons test (**A**) with p-values representing *p<0.05.

To address whether infectious particles were produced, we collected supernatant at 0, 1, and 2 days post-infection (dpi) from the same assay to quantify by plaque assay (**Fig 3C**). Corroborating our previous findings, infectious virus from CHIKV-IOL in the neutral pH 7.5 condition was not detectable, yet we found that even at a pH of 5.2, actively replicating CHIKV-IOL did not produce significant infectious virus. As a control for the effects of pH treatment, we found that MAYV reached high titers at 1 and 2 dpi as expected in both conditions. Overall, these results suggest that CHIKV-IOL is restricted in hCMECs at multiple steps in its life cycle, including both during endocytic entry and, based on the lack infectious particles, may also be restricted at egress.

### Prolonged ruxolitinib treatment of hCMECs rescues CHIKV-IOL infection and spread

It is known that endothelial cells produce a robust type-I interferon response, expressing a high level of interferon-stimulated genes (ISGs) at a basal level, and with strong induction after infection [23, 24]. Indeed, based on our previous co-localization analysis of CHIKV infection *in vivo*, we have seen an increase in the amount of actively replicating CHIKV-IOL colocalizing with cardiac endothelial cells in hearts from type I IFN receptor deficient mice as compared to WT C57BL/6 mice [10]. Therefore, we hypothesized that one mechanism of CHIKV-IOL restriction in human endothelial cells may be mediated by the type I interferon response. To evaluate this hypothesis, we used a JAK1/2 inhibitor, ruxolitinib (Rux), to inhibit the JAK/STAT pathway and downstream IFN signaling. We pretreated hCMECs with 5 μM Rux for 48 h, infected cells with CHIKV-IOL-ZsGreen with or without 5 μM Rux post-treatment and added fresh Rux daily for five days. After infection, we evaluated CHIKV-IOL-ZsGreen infection by high-content microscopy daily and at 5 dpi, quantified % infected as the number of ZsGreen positive cells over the number of total cells stained by DAPI (**Fig 4A and B**).

**Figure 4.**
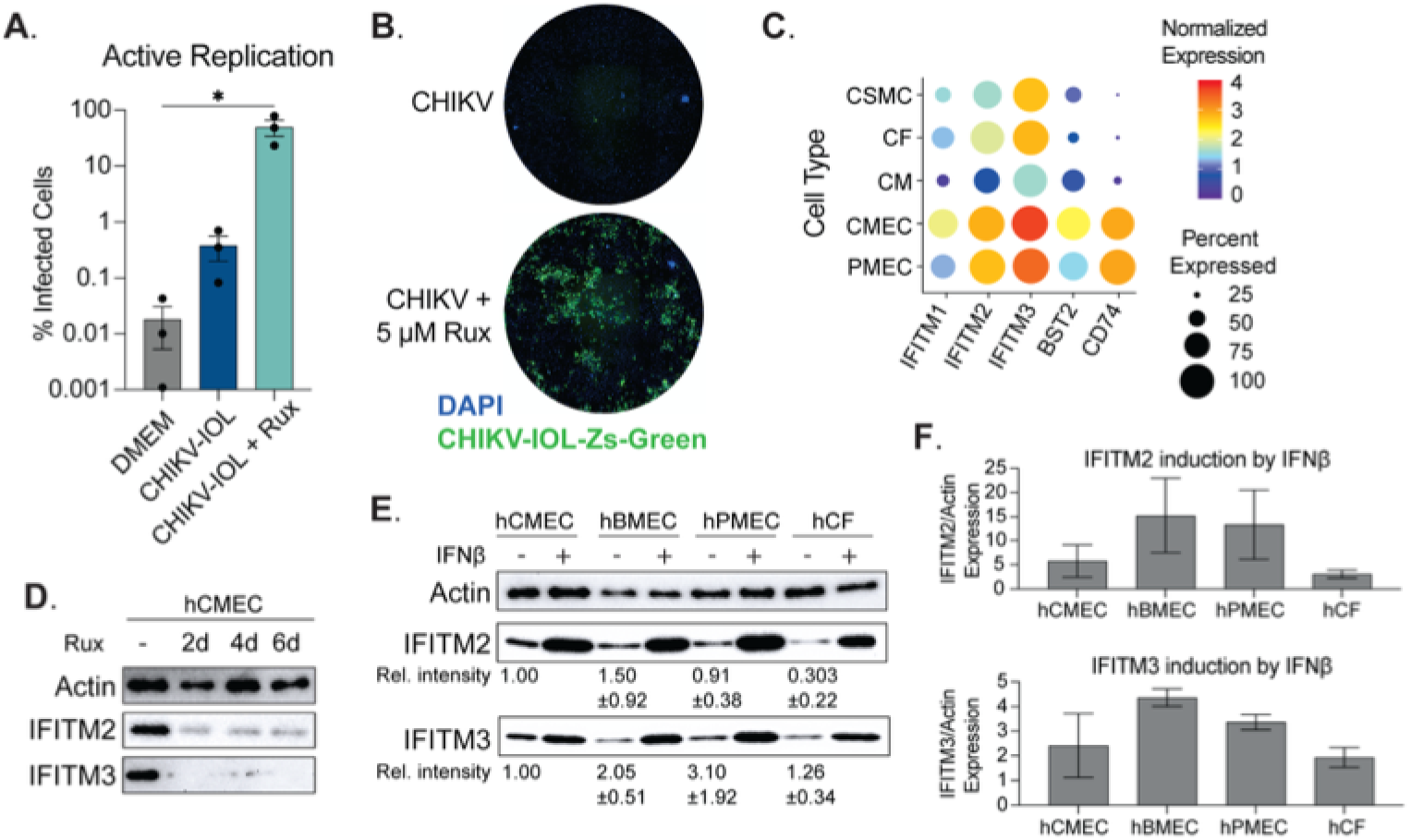
Primary cardiac endothelial cells are susceptible and permissive to CHIKV IOL in absence of JAK/STAT signaling. (**A**, **B**) HCMECs were pre-treated with 5 µM ruxolitinib or mock-treated for 2 days, then infected with CHIKV IOL ZsGreen at a MOI of 0.1 for 1 hour. Post-infection, cells were incubated with media supplemented with or without 5 µM ruxolitinib and fixed for high-content microscopy at 5 dpi. (**A**) Quantification of percent infected cells as measured by CHIKV-IOL-ZsGreen positive cells relative to number of total cells measured by DAPI staining. (**B**) Representative images from a CX7 high-content microscope showing cell nuclei stained with DAPI (blue channel) and CHIKV-IOL-ZsGreen infected cells (green channel). (**C**) Dot plot showing normalized expression levels of IFITM1, IFITM2, IFITM3, BST2, and CD74 in cardiac and endothelial cell types using the *Tabula Sapiens* dataset. (**D** and **E**) Representative images of western blots visualizing actin, IFITM2, and IFITM3 protein levels. (**D**) HCMECs were treated with 5 µM ruxolitinib for 2, 4, or 6 days or mock-treated. Cells were collected in laemmli buffer, proteins were separated by SDS-PAGE and visualized by immunoblotting. (**E, F**) HCMECs, hBMECs, hPMECs, and hCFs were treated with IFNβ or mock-treated for 24 hours. Cells were harvested and proteins analyzed as described above. (**E**) Values indicate relative intensity of expression as compared to hCMEC basal expression, with SD. (**F**) Quantified IFITM2 and IFITM3 expression after 24 hours IFNβ treatment. Data represents the mean of *n* = 3 independent trials in technical triplicate (**A, B**) with error bars showing the SEM and the limit of detection indicated by the gray shaded area. Western blots represent at least *n* = 2 independent trials (**D-F**). Statistical significance (**A**) was found by a Kruskal-Wallis test with multiple comparisons, with p-values representing *p<0.05.

In the untreated condition, we noted < 1% of all cells were positive for ZsGreen, comparable to the DMEM mock-infected condition. However, in the Rux treatment condition, an average of 49.9% of cells showed active replication across repetitions, with a statistically significant difference from the untreated control (**Fig 4A**). Notably, we did observe variations across technical replicates, though we consistently observed an increase in infection with Rux treatment, as well as spreading and CPE (**Fig 4B**). Interestingly, we did not observe ZsGreen expression until day 5 post infection (day 7 of Rux treatment), suggesting that it is the longterm Rux treatment of hCMECs that allows for viral replication. Given these results, we hypothesized that by treating cells for 7 days with Rux we could make the cells permissible for infection. However, under these conditions we saw no infection within 24 hours following 7 days of Rux treatment (data not shown), indicating that Rux, in combination with CHIKV infection, leads to replication overtime via unknown mechanisms. Overall, importantly, these results support that the restriction in endothelial cells is not mediated by the absence of receptor expression or replication co-factor. Moreover, these results suggest that restriction factors of CHIKV-IOL may be mediated by the JAK/STAT pathway.

### IFITM2 and IFITM3 are constitutively expressed in cardiac endothelial cells

Given that the restriction of CHIKV-IOL in hCMECs was at virus entry and egress and that inhibiting the JAK/STAT pathway could rescue CHIKV-IOL, we were interested in understanding the relative ISG expression in endothelial cells. Endothelial cells are known to constitutively express ISGs, which has been shown to play a role in protection from other viruses such as influenza virus and dengue virus [23–25]. Therefore, using the *Tabula Sapiens* single-cell transcriptomic atlas [26], we explored the constitutive expression of ISGs that were specific to human cardiac cells as well as hPMECs (**Fig 4C**). We first evaluated the expression of 66 known antiviral interferon-stimulated genes in human cardiac cell types [24, 25, 27, 28]. We selected for genes that were universally expressed in hCMECs, setting a cut-off value of expression in >70% of cells, yielding 5 genes of interest: IFITM1, IFITM2, IFITM3, BST2, and CD74 **(Fig 4C)**. Of these, all were significantly differentially expressed between hCMECs and other cardiac cell types and had an average Log_2_(Fold Change) > 2.7 between hCMECs and other cardiac cell types. Although scRNA data is not available for the brain within the *Tabula Sapiens* atlas, a comparison to hPMEC expression is also visualized. We found that similar to hCMECs, IFITM2 and IFITM3 were the highest basally expressed ISGs in hPMECs.

The IFITM family of proteins, of which IFITM2 and IFITM3 are expressed at a high basal level in hCMECs and hPMECs, have been shown to restrict several RNA viruses at endocytic entry, including influenza A, flaviviruses, and several alphaviruses [29–33]. To confirm the constitutive expression of these genes in our primary hCMECs and to evaluate the kinetics of Rux treatment, we treated cells with 5 uM Rux, replacing the treated media every 48 hours, and collected samples after 2, 4, and 6 days of treatment for IFITM2 and IFITM3 protein analysis. We confirmed the constitutive expression of both IFITM2 and IFITM3 in hCMECs, as well as the rapid and sustained reduction in IFITM2 and IFITM3 expression beginning at 2 days post-treatment **(Fig 4D)**. However, if constitutively expressed IFITM2 or IFITM3 were solely responsible for blocking CHIKV-IOL, we expected that the reduction noted at 2-day Rux pre-treatment should be sufficient to allow for the immediate recovery of CHIKV replication. Rather, we observed that it took up to 5 dpi for a notable number of ZsGreen cells to be observed by microscopy **(Fig 4A)**. These results again suggest that other mechanisms may be responsible for the rescue of CHIKV-IOL infection after Rux treatment of hCMECs.

To continue to explore the potential role of IFITM2 and IFITM3 in CHIKV-IOL infection specifically, we took advantage of the fact that CHIKV-IOL can infect hCFs, which also expresses IFITM3 at the RNA level (**Fig 4A**). We hypothesized that perhaps the protein level of IFITM2 and/or IFITM3 may be different between hCMECs and hCFs allowing for CHIKV-IOL infection, or that induction rather than basal expression may be involved. To test these hypotheses, we confirmed the constitutive expression of IFITM2 and IFITM3 across hCMECs, hPMECs, and hBMECs; cells which restricted CHIKV-IOL but not CHIKV-181/25 or MAYV. The relative intensity shown is calculated respective to hCMEC expression, with standard deviation indicated (**Fig 4E**). Interestingly, we confirmed that hCFs also expressed IFITM2 and IFITM3, although IFITM2 at a far lower level, as predicted by RNA (**Fig 4E and C**).

Finally, we wanted to know whether IFITM2 and IFITM3 could be stimulated to equal levels following interferon treatment. We stimulated all four cell types with IFNβ for 24 hours to evaluate the upregulation of IFITM2 and IFITM3 **(Fig 4E and F)**. Relative band intensity was quantified relative to actin control for each cell type (**Fig 4F**). We noted that IFITM2 and IFITM3 protein expression was upregulated highly across cell types in response to IFNβ, yet the magnitude of induction from baseline was different between cell lines suggesting differing antiviral responses. Interestingly, for both IFITM2 and IFITM3, the induction was lowest in hCFs and highest in hBMECs. However, it is important to consider than IFITM2 and IFITM3 are only two potential gene targets, used here as a proxy for IFN-signaling; indeed, other ISGs with distinct responses may be playing a role.

Overall, these results support the literature suggesting that endothelial cells not only have a high basal interferon response but also produce a robust innate immune response after infection as compared to other cell types. We found that CHIKV-IOL infection can be recovered in the presence of a JAK/STAT inhibitor, yet the exact mechanism remains elusive. Future studies will be required to understand the multiple levels of restriction in endothelial cells.

### MAYV does not induce a potent type I interferon response during infection of cardiac endothelial cells

Given that MAYV was able to infect nearly the entire endothelial cell monolayer while CHIKV-181/25 replicated in patches (**Fig 2C**), we hypothesized that MAYV may antagonize the interferon response leading to spread. Therefore, we explored the type-I IFN response in hCMECs after CHIKV and MAYV infection. To compare virus infection accurately, we generated all virus stocks in low-passage BHK-21 cells and purified them via ultracentrifugation over a 20% sucrose cushion. We infected hCMECs with purified CHIKV-IOL, CHIKV-181/25, or MAYV, as well as with corresponding heat-inactivated (HI) controls, or mock-infected (DMEM) controls at a MOI of 1. At 0 and 24 hpi, we collected supernatants to quantify infectious particles by plaque assay (**Fig 5A**) and then collected the cell monolayer for viral and cellular RNA analysis by RT-qPCR (**Fig 5B and C**). In line with all previous experiments, CHIKV-IOL infectious particles were not detectable at 24 hpi, while CHIKV-181/25 and MAYV productively infected hCMECs (**Fig 5A**). At 24 hpi, MAYV and CHIKV-181/25 titers were significantly different from CHIKV-IOL titers, further supporting our previous results (**Fig 2A**). To confirm viral replication, we also quantified CHIKV RNA by Taqman RT-qPCR (**Fig 5B**). Although both CHIKV-IOL and CHIKV-181/25 started at comparable levels of RNA at 0 hpi, CHIKV-181/25 RNA significantly increased corresponding with viral replication. Indeed, supporting differences in infectious particle production, a statistically significant difference in viral RNA between both strains was found at 24 hpi.

**Figure 5.**
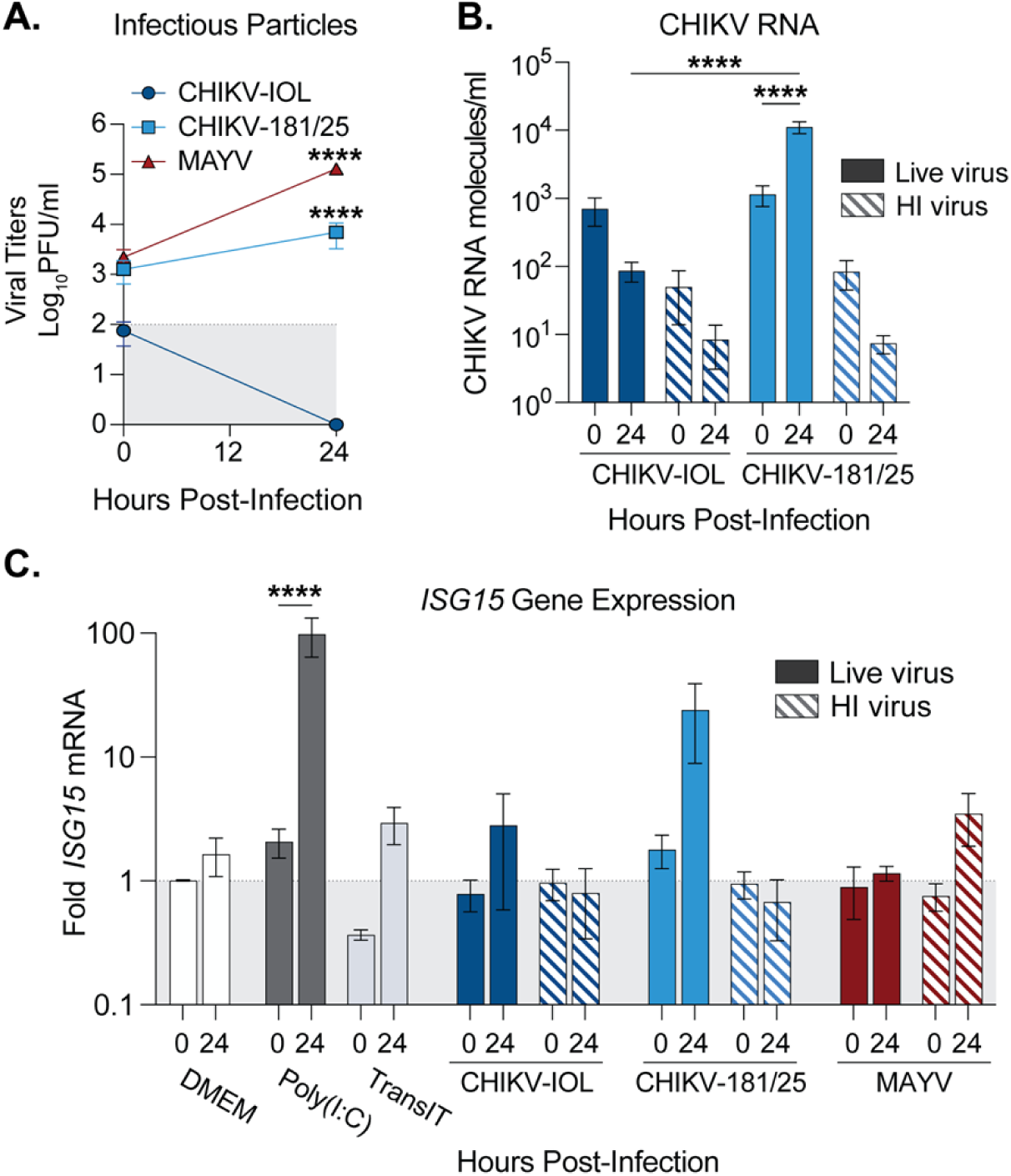
MAYV infection attenuates the type-I IFN response in cardiac endothelial cells. (**A-C**) hCMECs were either pre-treated with Poly(I:C) or mock-transfected for 2 hours and then mock-infected with DMEM, or infected with CHIKV-IOL, CHIKV-181/25, MAYV, corresponding heat-inactivated controls (HI), or mock-infected (DMEM) at a MOI of 1 for 1 hour. Supernatants and cell monolayers were collected at 0 and 24 hpi. (**A**) Viral titers were quantified by plaque assay and (**B**) intracellular viral genomes were quantified by RT-qPCR. (**C**) *ISG15* gene expression quantified relative to DMEM for each timepoint by RT-qPCR. Data represents the mean of *n* = 3 independent trials in technical duplicate with error bars representing the SEM and the limit of detection indicated by the gray shaded area. Statistical significance was found by (**A - C**) two-way ANOVA with multiple comparisons, with statistics in (**A**) representing comparisons to CHIKV-IOL at 24 hpi. P-values represent ****p<0.0001.

To address the innate immune response after infection, we next evaluated *ISG15* host gene expression by RT-qPCR. *ISG15* was selected as a potent interferon-stimulated gene to evaluate ISG induction. In the same experiment as above, wells were also pre-treated with Poly(I:C) as a positive control, or mock-transfected (TransIT) for 3 hours, in addition to the cells infected with live virus or HI controls. As expected, at 24 hpi, in cells treated with Poly(I:C), there was nearly a 100-fold increase in *ISG15* mRNA as compared to mock-infected cells (DMEM) or transfection reagent alone (**Fig 5C**). Given the difference in infectious particles and RNA between CHIKV-IOL and CHIKV-181/25, it aligns that CHIKV-181/25 infection produces a marked increase in *ISG15* at 24 hours, while the non-replicating CHIKV-IOL does not significantly induce *ISG15* above the control (**Fig 5C**). Additionally, an *ISG15* increase was not noted in the CHIKV-181/25 HI control, suggesting that *ISG15* production was a result of actively replicating CHIKV-181/25 virus. Interestingly, despite reaching the highest viral titers, MAYV did not produce any *ISG15* response above baseline at 24 hours (**Fig 5C**). These results suggest that hCMECs respond differently to different alphaviruses and that MAYV, while replicating to high levels, does not induce a potent IFN response in human endothelial cells.

## Discussion

Arboviruses include various human pathogens of significance to public health, many of which are poorly surveilled and understudied. Specifically, the ability for emerging arboviruses and alphaviruses to infect different cell types and tissues is incompletely understood. Here, we used human primary cardiac cells to characterize the ability for different arboviruses to directly infect human cardiac cell types *in vitro*. We find that ZIKV, RVFV, LACV, and MAYV can infect multiple human primary cardiac cell types. These findings are particularly interesting in the case of ZIKV, which has reported cardiovascular manifestations in adult humans, but has had limited investigation *in vitro* as to the mechanism of pathogenesis. Additionally, LACV, an orthobunyavirus, was recently found to replicate in the heart of infected mice at 3 dpi [34], while its tropism for cardiac cells *in vitro* has not yet been explored. Finally, given the cardiac pathogenesis characterized after CHIKV infection *in vivo* [10], MAYV’s broad tropism for cardiac cells is particularly important.

In our prior work, we found that the CHIKV-IOL strain specifically targeted hCFs, while infection was restricted by hCMECs [10], a contrast from the tropism we characterized with MAYV *in vitro*. Beyond the heart, microvascular endothelial cells are important for lining and protecting the vasculature. Considering that arboviruses are found in high titers in circulation, the interactions between these viruses and endothelial cells may have significant implications for viral dissemination in multiple organs. Indeed, differences in tropism in hCMECs between CHIKV and MAYV were replicated in hPMECs and hBMECs. Characterizing this difference further, we found this restriction to be specific to the CHIKV-IOL strain across multiple endothelial cell types. Moreover, restriction appears to be occurring at both endocytic entry and at viral egress, and in hCMECs, downstream of the JAK/STAT pathway.

Although further work is needed to characterize the specific restriction factors playing a role, one hypothesis is that constitutively expressed and/or IFN inducible genes, such as the IFITM family, BST2, or others, are restricting CHIKV-IOL at multiple steps in the life cycle. Among alphaviruses, IFITM2 and IFITM3, and to a lesser extent IFITM1, have been found to restrict SINV and SFV infection in A549 cells [35]. In addition, IFITM3 has been implicated in CHIKV pathogenesis *in vitro* and *in vivo*, as well as in SFV, ONNV, VEEV, and SINV infection in mouse embryonic fibroblasts [33]. Additionally, both CHIKV and MAYV were found to be restricted by IFITM3 in HEK293T cells and HeLa cells [36]. We also identified BST2 (Tetherin) in our search, an ISG known to broadly prevent enveloped virus budding [37]. Among alphaviruses, BST2 has been shown to prevent SFV and CHIKV VLP release [38]. Finally, CD74, a major histocompatibility complex class II chaperone, has been shown to inhibit the viral entry of both Ebola and SARS-CoV-2 [39]. An important alternative hypothesis may be that Ruxolitinib treatment, as a JAK1/2 inhibitor, may have unknown off-target affects from the type I interferon pathway that are playing a role in CHIKV-IOL restriction. Nonetheless, future work to address these restriction factors is important. Primary endothelial cells are notoriously difficult to transfect, rendering siRNA knockdowns of target genes challenging. Therefore, future work might utilize alternative and unbiased screening techniques to identify factors specific to endothelial cells that restrict CHIKV-IOL in a finely tuned manner.

In addition to cellular restriction factors, these studies highlight the presence of viral factors that contribute to infection and the innate immune response. Our findings suggest that MAYV may be bypassing or antagonizing the innate immune response in endothelial cells, allowing the virus to replicate more efficiently than other alphaviruses. MAYV is an emerging alphavirus that, although less well studied than CHIKV, has the potential to adapt to urban environments and warrants further investigation [40, 41]. Indeed, a recent study noted differences between Mayaro virus and Una virus (UNAV), another related alphavirus, during the infection of immortalized human brain microvascular endothelial cells [42]. Similar to our findings, the authors found that MAYV infected these cells, albeit to a low titer, while UNAV infectious titers nor proteins were detectable. In addition to interactions with specific ISGs, it is possible that MAYV may be using an alternative receptor or alternative entry route that escapes these restrictions or evades sensing pathways. Moreover, although CHIKV has been shown to infect endothelial cells in culture, these studies use other CHIKV strains such as the lab-adapted Ross strain or found mixed results depending on the endothelial cell type [13, 15]. Therefore, our study highlights the importance of virus strain, such as those which are not lab adapted, and relevant cell types, such as primary models, as restriction appears highly specific to the CHIKV-IOL strain used. Further work is critical to interrogate the factors that enable MAYV to bypass this restriction, but also to identify the genetic elements that render CHIKV-IOL sensitive.

Taken together, our findings suggest various repercussions for viral pathogenesis in a living model. However, there are limitations to these *in vitro* models. First, they require virus to be placed directly on a cell monolayer, limiting the complexity as compared to an *in vivo* model; differences in cell tropism between *in vitro* and *in vivo* experiments may be mediated by factors such as cell geography, viral dissemination and changes in primary cell phenotype in an *in vitro* environment. One limitation of this study is that for the primary cells, we only analyzed cells from one donor. While these results were consistent across endothelial cells, differences between individuals may impact infection. Despite these limitations, using physiologically relevant models, we found that multiple arboviruses from diverse families can infect cardiac cells *in vitro.* Future *in vitro* and *in vivo* studies with these emerging viruses will be critical to understand viral tropism in the heart and the cardiovascular system.

Overall, this work characterizes the differential restriction of alphaviruses across endothelial cell types, finding CHIKV to be restricted in a strain-specific manner. Importantly, the related and emerging MAYV produces high viral titers across multiple human endothelial cell types, which warrants further investigation. Although further work is needed to explore the specific host factors involved in restriction, and the viral genomic elements that render MAYV resistant to restriction, these findings suggest different mechanisms of pathogenesis across alphaviruses.

## Acknowledgements

We thank all members of the Stapleford lab for their insight on this work. We thank Dr. Meike Dittmann at the NYU Grossman School of Medicine for use of the CX7 CellInsight microscope, and for gifting poly(I:C) reagent. We thank Drs. Gonzalo Moratorio and Alvaro Fajardo at the Institut Pasteur de Montevideo for gifting MAYV, as well as Dr. Ana Rodriguez and Kelly Crotty at the NYU Grossman School of Medicine for gifting hBMECs and hPMECs. Finally, we thank Drs. Ludo Desvignes and Dominick Papandrea for use of the NYU Grossman School of Medicine ABSL3 facility. This work was supported by funding from the NYUGSoM Start up, the American Heart Association Postdoctoral Fellowship (19-A0-00-1003686) (M.G.N.), and NIH/NIAID R01 AI162774-01A1 (K.A.S.).

## Competing Interests

The authors declare no competing interests.

## Notes

### Competing Interest Statement

The authors have declared no competing interest.

